# Parabrachial *Calca* neurons influence aversive and appetitive taste function

**DOI:** 10.1101/2025.02.28.640892

**Authors:** Christian H. Lemon, Jinrong Li, Md Sams Sazzad Ali, Neville M. Ngum, Kyle T. Zumpano, Catori J. Roberts

**Affiliations:** School of Biological Sciences, University of Oklahoma, 101 David L. Boren Blvd., Norman, OK 73019 USA

## Abstract

The parabrachial (PB) nucleus participates in taste processing and integration with other senses. PB neurons that express the *Calca* gene support sensory-integrative responses, albeit only limited data have addressed their influence on taste. Here we studied how chemogenetic dampening of PB-*Calca* neurons impacted mouse orosensory preferences for diverse taste stimuli in brief-access fluid exposure tests, which capture oral sensory/tongue control of licking behavior. Intracranial delivery of Cre-dependent viruses in female and male *Calca*^Cre/+^ mice induced expression of the inhibitory designer receptor hM4Di:mCherry (hM4Di mice) or fluorophore mCherry alone (mCherry mice) in PB-*Calca* neurons. Several weeks later, hM4Di and mCherry mice entered brief-access tests where they could lick taste solutions on discrete seconds-long trials. Stimuli included the behaviorally avoided, but functionally different, bitter taste stimuli quinine (0 [water], 0.1, 0.3, and 1.0 mM) and cycloheximide (0, 0.001, 0.003, and 0.01 mM), and the appetitive sugar sucrose (0, 100, 300, 500, and 1000 mM). Both hM4Di and mCherry mice received the hM4Di ligand clozapine-N-oxide (CNO, 5 mg/kg, i.p.) prior to daily tests performed by blinded experimenters. With CNO, hM4Di mice displayed greater average licking (i.e., less avoidance) of quinine (p < 0.05), but not cycloheximide (p > 0.3), than mCherry mice, implying PB-*Calca* neurons variably influence orosensory responses across bitter stimuli. Moreover, male hM4Di mice selectively showed reduced mean licking preferences for sucrose under CNO (p < 0.05). These data suggest that PB-*Calca* neurons participate in both aversive and appetitive taste-guided behaviors, with their role in appetitive taste dependent on sex.

## Introduction

The sense of taste participates in the recognition of nutrient and toxin chemicals in potential food sources. In mammals, these chemicals engage diverse families of taste receptor proteins expressed by receptor cells comprising taste buds on lingual and intraoral surfaces (Roper and Chaudhari, 2017; Kinnamon and Finger, 2019). Gustatory neural information is relayed to the brain by cranial nerve fibers that terminate in the rostral nucleus of the solitary tract (rNTS) in the medulla (Pfaffmann et al., 1961; Halpern and Nelson, 1965). In rodents, the rNTS houses neurons that project ascending gustatory signals to the parabrachial (PB) nucleus of the pons (Norgren, 1974). The PB area is implicated with multiple homeostatic and sensory roles and maintains neurons with axons that reach diverse brain regions associated with affect, including the amygdala and bed nucleus of the stria terminalis (Norgren, 1976; Saper and Loewy, 1980; Fulwiler and Saper, 1984; Gauriau and Bernard, 2002; Chiang et al., 2019).

A growing literature has described that both the rNTS and PB nucleus are composed of heterogeneous genetic cell types that influence specific aspects of taste processing. For instance, the rNTS is populated by γ-aminobutyric acid-positive inhibitory neurons evidenced to modulate the intensity of neural and behavioral responses to tastes (Travers et al., 2022). Other rNTS neuron types were also associated with taste processing (Zhang et al., 2019; Jin et al., 2021). In the PB area, neurons marked by the transcription factor Satb2 were implicated in taste function (Fu et al., 2019; Jarvie et al., 2021), with some evidence to suggest that PB neurons expressing *Calca* may participate in responses to aversive tastes (Jarvie et al., 2021; Kang et al., 2022; Kim et al., 2024b). Notably, the *Calca* gene is a marker of PB neurons that express the neuropeptide CGRP (calcitonin gene-related peptide) and are implicated for roles in homeostasis and protection, including regulation of appetite and mediating pain-related responses (Carter et al., 2013; Campos et al., 2018; Palmiter, 2018).

*Calca* neurons densely populate lateral regions of the PB area identified, using neurophysiology and optogenetics, to contain taste-active cells that receive convergent input from ascending trigeminal somatosensory circuits (Li and Lemon, 2019; Li et al., 2022). Some of these taste-integrative neurons respond to both gustatory and intraoral trigeminal stimuli, activating to chemesthetic and thermal nociceptive inputs (trigeminal) and the bitter tastants quinine and cycloheximide (Li and Lemon, 2019; Li et al., 2022). Quinine and cycloheximide are ecologically and functionally diverse bitter-tasting chemicals/toxins that elicit concentration-dependent reductions in fluid licking (i.e., avoidance) in rodents (Boughter et al., 2005; Hettinger et al., 2007; Travers and Geran, 2009; Wilson et al., 2012). Based on their location, PB taste-integrative neurons may include cells marked by the *Calca* gene, although the role of *Calca* neurons in orosensory responses to diverse bitter tastes is unknown.

Here, we used Cre-directed chemogenetics to study how temporary dampening of activity in PB-*Calca* neurons affected licking-avoidance behaviors that mice show towards the bitter taste stimuli quinine and cycloheximide. We also examined how chemogenetic suppression of PB-*Calca* cells influenced mouse licking preferences for the preferred sugar sucrose. These studies were accomplished using a large number of experimental and control mice of both sexes tested by blinded experimenters in brief-access lickometry tests. Brief-access tests monitor rodent licking responses to fluids on short, seconds-long trials, which captures oral sensory guidance of ingestive preference while mitigating post-ingestive influences (Smith, 2001; Boughter et al., 2002).

Analyses revealed that during chemogenetic suppression targeted to PB-*Calca* neurons, mice increased the number of licks they emitted to quinine (i.e., showed reduced quinine avoidance), with the magnitude of increase variable across animals. In contrast, licking responses to cycloheximide were unaffected. These findings suggest that *Calca* cells differentially influence orosensory avoidance behavior across diverse bitter taste stimuli. Furthermore, suppression of PB-*Calca* neurons reduced mouse orosensory licking preferences for appetitive sucrose, with reductions in sucrose licking showing significant sex dependence and emerging only in males. That perturbation of PB-*Calca* neurons affected both avoided (quinine) and preferred (sucrose) tastes in our studies may relate to recent data on an ability of this cell class to bidirectionally encode appetitive and aversive stimuli (Kim et al., 2024a). Lastly, our results concerning participation of PB-*Calca* neurons with sucrose taste preference also represent a sex-specific brain cell-type effect on gustatory behavior.

## Methods

### Mice

All experiments and procedures were approved by the Institutional Animal Care and Use Committee at the University of Oklahoma and followed the *Guide for the Care and Use of Laboratory Animals* by the National Research Council. All studies herein used adult female and male *Calca*^Cre/+^ mice. These mice were generated by crossing homozygous *Calca*^Cre^ mice (Strain #033168, The Jackson Laboratory [JAX], Bar Harbor, ME) (Carter et al., 2013) with C57BL/6J mice (Strain #000664, JAX). A total of 88 *Calca*^Cre/+^mice were surgically prepared and tested in these studies. At the onset of behavioral testing, study mice were 34.0 (mean) ± 12.8 (std. dev.) weeks of age. On average, females (*n* = 47) weighed 23.4 ± 3.0 g whereas males (*n* = 41) weighed 31.5 ± 3.9 g.

Before surgery, mice were group housed in disposable cages with *ad libitum* access to standard mouse chow and filtered water. After surgery, each study mouse was singly housed in a disposable cage, with water access regulated during behavioral studies, as below. Mouse cages were housed in an individually ventilated (HEPA filtered) racking system (Innovive, San Diego, CA) located in a climate-controlled room maintained on a 12 hr light/dark cycle. Individual cages were prepared with standard bedding and enrichment materials (e.g., paper huts, carboard tubes). Leading up to training for behavioral tests, bedding was changed by husbandry staff on a regular schedule. During experiments, bedding was changed by experimenters as needed to avoid housing disruptions.

### Surgery and bilateral intracranial microinjections

An intracranial microinjection procedure was used for viral delivery of fluorophore and DREADD (designer receptors exclusively activated by designer drugs) proteins to PB neurons. Mice were divided into experimental and control groups depending on the type of virus received. Experimental *Calca*^Cre/+^ mice (hereafter referred to as hM4Di mice) received bilateral microinjections of a Cre-dependent virus enabling cellular expression of the inhibitory Gi-DREADD hM4Di paired with the fluorophore mCherry (AAV1-hSyn-DIO-hM4D(Gi)-mCherry, #44362, Addgene, Watertown, MA). When expressed in neurons and stimulated with clozapine N-oxide (CNO), hM4Di reduces neural firing (Urban and Roth, 2015; Roth, 2016), including in PB-*Calca* neurons (Carter et al., 2013). Control *Calca*^Cre/+^ mice (hereafter known as mCherry mice) received bilateral microinjections of a virus supporting Cre-activated expression of mCherry alone (AAV1-hSyn-DIO-mCherry, #50459, Addgene).

Over the course of these studies, groups of about 9 mice, on average, were surgically prepared (one or two per day) and then, following recovery and a waiting period, tested together in behavioral studies as a squad. Ten squads were examined. Each squad included female and male mCherry mice and female and male hM4Di mice to avoid temporal confounds with testing different mouse groups/sexes.

All surgical tools were sterilized prior to use. The stereotaxic device (Model 1900 Stereotaxic Alignment System, Kopf Instruments, Tujunga, CA) and surrounding workstation/countertop area were cleaned to prepare a sterile field. Approximately 30 min prior to surgery, mice received an injection of the antibiotic gentamicin (5 mg/kg, s.c.) to ward off the potential for infection. Mice were then placed in a clear plexiglass rodent induction chamber and anesthetized with ∼3% isoflurane in oxygen, delivered at about 1 L/min. Once anesthetized, the scalp was shaved free of fur and mice were transferred to the stereotaxic head frame with their snout positioned in a gas anesthesia nose cone with incisor bar. Anesthesia was maintained by administrating 1-3% isoflurane in oxygen, delivered at about 0.6 to 1 L/min. The mouse was positioned atop a feedback-controlled heating pad maintained at 37°C. Lubricating eye (ophthalmic) ointment was applied to both eyes.

Anesthesia depth was monitored by absence of eye blink, the absence of foot withdrawal following heavy pinch, a lack of startle from tail pinch or by monitoring for any other overt signs of response to physical stimuli. Once surgical-level anesthesia was ensured, an antiseptic (70% ethanol) followed by antibiotic (betadine) were topically applied to the bare scalp. A midline incision was made to expose the cranium. The skull was then brought into final alignment and leveled using stereotaxic measurements made from cranial fissures. Coordinates for targeting the microinjection needle tip to the PB area were: 4.9 to 5.3 mm caudal of Bregma, 1.0 to 1.3 mm lateral, and about 2.6 mm ventral from the brain surface. These coordinates were obtained from published (Franklin and Paxinos, 2008) and online (Allen Brain Atlas) sources.

A sterilized drill bit was used to make a small craniotomy at the targeted location on the skull. A glass Hamilton microsyringe coupled to a sterilized 33-gauge beveled needle was then positioned perpendicular to the skull in a syringe pump (Micro 4 MicroSyringe pump, World Precision Instruments, Sarasota, FL) coupled to the stereotaxic device. Based on coordinates, the needle tip was then slowly and precisely lowered into brain tissue to reach the PB area using the fine-control stereotaxic manipulator arms, with tip position tracked using a digital readout.

After waiting for about 10 minutes to allow brain tissue to recover from needle insertion, 0.5 µL of the virus was ejected from the syringe needle tip at a pump-controlled rate of 0.05 µL/min. Once the injection was complete, the needle remained in place for an additional 5 to 10 min before being slowly withdrawn dorsally from the brain and skull using fine controls on the stereotaxic device. The craniotomy and microinjection procedures were repeated for the contralateral PB nucleus.

Prior to closing the incision site, bone wax was applied to the craniotomies to seal them. One drop of the long-lasting local anesthetic bupivacaine was applied to the skin and periosteum surrounding the craniotomy. The scalp incision was closed by silk suture. Mice were then removed from the stereotaxic device, hydrated with 0.5 mL lactated ringers (s.c.), and administered buprenex (0.05 to 0.2 mg/kg, i.p.) for management of potential discomfort or pain.

Once mobile after surgery, each mouse was singly housed in a small disposable cage in our colony rack and monitored daily until it fully recovered from the procedure. Mice typically showed normal ambulatory behavior the day after surgery. Sutures disappeared after several days. No mice required additional analgesics during recovery. Mice entering behavioral tests > 8 weeks following surgery.

### Experimenter blinding

Prior to behavioral testing, a lab confederate matched a unique alphanumeric code to each study mouse and randomized their cage locations in the colony housing rack before training commenced. The codes were used to label mice in lieu of all other identifying mouse information, which blinded the experimenters handing and running mice, including administering CNO, to mouse DREADD group (i.e., hM4Di or mCherry). All mice had black coats and indistinguishable outward appearances, facilitating blinding. Blinding was used to collect behavioral data across all mouse training and test sessions and for scoring brain tissue fluorescence, as below.

### Water restriction schedule

An overnight water restriction schedule was imposed on study mice the day before their lickometer training commenced. This schedule aimed to motivate mice to lick fluids offered during brief-access fluid exposure sessions. To do this, each water bottle was removed from the home cage and replaced with a marble-weighted bottle with no water, which blocked the bottle access hole on the cage top. With exceptions noted below, water restriction conditions continued through testing, with mice consuming their daily fluids during training and test sessions in lickometers. During all study phases, mice were given 1-hr access to water in their home cage if their daily measured body weight fell below about 80% of their baseline weight. Food was always freely available to mice in their single housing cages.

### Lickometer apparatus

Behavioral tests were carried out using a “Davis Rig” contact lickometer (DiLog Instruments, Tallahassee, Florida; Med Associates, St. Albans, VT). This computer-controlled device can record licking responses made by a mouse to different fluids offered on sequential, seconds-long trials while randomizing the order of fluid presentation within one test session. The short length and limited number of fluid trials offered during brief-access tests mitigates post-ingestive influences on licking responses, capturing ingestive/licking behavior driven by initial oral sensation (Smith, 2001; Boughter et al., 2002). For our studies, several Davis Rigs were used in parallel and in series, with mouse test order and rig assignment randomly determined daily. Mouse licking responses to temperature-controlled fluids were recorded using a custom-modified “thermolickometer” Davis Rig we developed (Li et al., 2024).

### Lickometer training

Overnight water-restricted mice were individually trained to receive fluids in a Davis Rig over four days. On training days 1 and 2, mice were allowed free access to one sipper tube filled with room temperature water for 30 min to habituate them to the apparatus. Days 3 and 4 familiarized mice with the brief-access fluid exposure procedure, offering them sipper tubes of room temperature water over 20, 10-sec access trials. Once a tube was presented, mice were allowed 30 sec to make a lick, which started the trial. Zero licks were recorded if no licks were made after 30 sec. Inter-trial intervals were 7.5 or 10 sec. After completing the last day of training, mice were returned to their single-housing cage and given *ad lib* access to water for approximately two days, prior to beginning testing.

### CNO administration

Brief-access fluid exposure tests were performed with, and in some cases without, administration of the hM4Di agonist CNO (Sigma-Aldrich). On test days conducted under CNO influence, all mCherry and hM4Di mice in the squad received an injection (i.p.) of CNO (5 mg/kg) about 30 min prior to the start of their brief-access session. The experimental blinding procedure described above prevented experimenters from knowing whether mice belonged to the mCherry or hM4Di group. CNO dosing follows published recommendations for mice (Jendryka et al., 2019). CNO (5 mg vial) was dissolved in 50 µL of DMSO and then transferred to sterilized saline (up to 10 mL).

Following Cre-directed viral transfection, PB-*Calca* neurons in hM4Di mice expressed the mCherry fluorophore and hM4Di – an inhibitory DREADD engaged by CNO. Thus, administering CNO to hM4Di mice supported temporary dampening of activity in PB-*Calca* neurons during brief-access tests. Receipt of CNO by mCherry mice, where PB-*Calca* neurons expressed only mCherry, controlled for off-target effects of this ligand (Mahler and Aston-Jones, 2018; Jendryka et al., 2019).

### Brief-access ffuid exposure tests

After an approximate two-day break with *ad lib* access to water following training, mice returned to an overnight water-restriction schedule and entered brief-access fluid exposure tests with orosensory stimuli conducted in the lickometers. All chemical stimuli were dissolved in purified water (hereafter referred to as water). Four different stimuli, including innocuous thermal and hedonic taste stimuli, were tested as follows:

1. *Temperature-controlled water*. Overnight water-restricted mice were proffered water precisely held at 15°C (mild cooling) or 35°C (innocuous warmth) over a series 5 sec brief-access trials arranged in random interleaved order. Each water temperature was offered on 15 trials (30 trials total per session; mice were allowed up to 75 sec of licking access to each water temperature per day). This test was administered over 4 consecutive days, with each mouse receiving an injection of CNO about 30 minutes prior to starting daily testing.
2. *Quinine*. Three different concentrations of the ionic bitter taste stimulus quinine-HCl (0.1, 0.3, and 1.0 mM) and water (0 mM) were proffered to overnight water-restricted mice on 10-sec trials, with fluid order randomized within each of 5 contiguous trial blocks (20 trials total; mice were allowed up to 50 sec of licking access to each concentration per day). Quinine solutions were presented at room temperature. This test was repeated over 4 consecutive days, with CNO administered to mice each day about 30 min prior to the start of testing.
3. *Cycloheximide*. Overnight water-restricted mice were presented three different concentrations of the non-ionic bitter taste stimulus cycloheximide (0.001, 0.003, and 0.01 mM) and water (0 mM) on 10-sec trials, with fluid order randomized within each of 5 contiguous trial blocks (20 trials total; mice were allowed up to 50 sec of licking access to each concentration per day). Cycloheximide solutions were offered at room temperature. These tests were conducted over 4 consecutive days under CNO, with CNO administered about 30 min prior to testing each day.
4. *Sucrose*. Brief-access fluid exposure tests with sucrose were performed in two phases. In phase 1, overnight water-restricted mice were proffered four concentrations of sucrose (100, 300, 500, and 1000 mM) and water (0 mM) at room temperature in a brief access setting for 2 days, without CNO. Trials were 5-sec long with stimulus order randomized over 8 blocks (40 trials total; mice were allowed up to 40 sec of licking access to each concentration per day). Phase 1 was a training session to allow mice to learn that the sipper tubes offered sucrose – a strongly preferred taste stimulus.

On completion of phase 1, mice were given full access to water in their home cage and entered phase 2, which began the next day. Phase 2 monitored brief-access licking responses to sucrose in water-replete (not thirsty) mice with continuous access to water in their home cage. Thus, mice responded to the hedonic features of sucrose taste in the absence of thirst motivation. During each test session, mice were proffered 4 different concentrations of sucrose (100, 300, 500, and 1000 mM) and water (0 mM) on discrete 5-sec trials, with fluid order randomized within each of 8 contiguous trial blocks (40 trials total; mice were allowed up to 40 sec of licking access to each concentration per day). Solutions were offered at room temperature. Tests were conducted across 4 sequential days, with CNO administered about 30 minutes prior to each daily session.

A subset of mice entered brief-access tests with sucrose solutions conducted as in phase 2, but without daily administration of CNO. This no-CNO condition allowed for analysis of orosensory responses to sucrose captured with and without CNO-induced perturbation of PB-Calca neurons. In most cases, the 4-day no-CNO tests with sucrose began the day after the 4-day with-CNO tests completed.

Concentration series for each taste stimulus followed published literature and elicited mild, moderate, to strong effects on licking behavior with increasing values. The tested water temperatures stimulate (15°C) or reduce (35°C) licking in brief-access tests in mice (Li et al., 2024).

Most mice were tested with multiple stimuli, with a break of at least 4 days separating tests with different solutions. All mice within a squad experienced the same stimuli series and test conditions. Mice received *ad lib* access to water in their single-housing cage during break periods. No mice underwent all stimulus tests; sample sizes for specific tests are detailed in the results.

### Fluorescence microscopy of brain tissue

At the conclusion of behavioral testing, mice were euthanized by an overdose of sodium pentobarbital (≥130 mg/kg, i.p.) and underwent transcardial perfusion with 0.9% NaCl followed by 4% paraformaldehyde and 3% sucrose dissolved in 0.1 M phosphate buffer. Brains were removed and stored in 4% paraformaldehyde and 20% sucrose dissolved in 0.1 M phosphate buffer refrigerated to ∼4°C. A sliding microtome (SM2010R, Leica) was used to cut coronal sections (20 µm) through midbrain/brain stem tissue that included the PB area. A fluorescence microscope (Axio Scope.A1, Carl Zeiss Microscopy) and camera system (Axiocam 305 with ZEN software, Zeiss) were used by an experimenter blind to mouse group to inspect sections for mCherry (mCherry alone and hM4Di:mCherry) fluorescence labeling of neurons in the PB region.

### Data analysis and statistics

The number of licks mice emitted on each brief-access trial were calculated by adding 1 to the number of inter-lick intervals that were >50 msec. This criterion filtered any erroneously recorded phantom/noise activity (e.g., Ellingson et al., 2009; Lemon et al., 2019; Li et al., 2024). For each stimulus/concentration, the mean number of licks that a mouse emitted across all trials and test days was calculated by applying a 20% trimmed mean to the calculated lick counts, ignoring trials where no sampling occurred (i.e., zero licks). The trimmed mean accommodated outliers by computing average licks after dropping the lower 20% and upper 20% of lick counts considered. As a convenience, mean licks per trial is referred to as “licks” going forward.

Licks to the aversive bitter taste stimuli quinine and cycloheximide were also expressed as lick scores, calculated for each mouse as licks to each stimulus/concentration minus licks to water. Licks to water was obtained from analysis of water trials randomly proffered with the stimulus series during testing (i.e., the 0 mM trials). Lick scores standardized responses to aversive tastes as decreases in licking from water responding.

Licks to appetitive sucrose were analyzed using lick ratios, calculated for each mouse by dividing their licks made to each sucrose concentration by their mean primary lick rate. Mean primary lick rates were based on the mean primary inter-lick interval (MPI) to water observed for each mouse during the first 5 trials of the brief-access training days, using inter-lick intervals ≥50 and ≤160 msec. This window reflects the primary component of the inter-lick interval distribution, which includes most intervals/licks a mouse typically emits (Boughter et al., 2012). The reciprocal of the MPI was taken to derive the mean primary lick rate (licks/sec), which was then scaled to the length of the stimulus trial under consideration (licks/5 sec) to estimate the number of licks that each mouse could theoretically achieve in that timeframe with constant licking. Sucrose lick ratios that approached 1 indicated near maximal licking; those that approached 0 reflected minimal licking. Our calculation of lick ratios is appropriate to standardize orosensory responses and gauge avidity to appetitive taste solutions like sucrose, which evokes increases in responding with rising concentration in the absence or reduction of thirst motivation (e.g., Glendinning et al., 2002; Dotson and Spector, 2004; Ellingson et al., 2009).

Latency to first lick from shutter opening was also considered for each stimulus to gauge the potential influence of olfactory/vapor cues on licking behaviors. Olfactory influences on mouse licking in brief-access assays can appear as a systematic change in latency with change in stimulus concentration (Boughter et al., 2002; Glendinning et al., 2002). For individual stimuli and mice, latency to first lick was the average (20% trimmed mean) of the latencies collected across all sampled trials.

Statistical analyses coupled inferential with estimation statistics to study the significance and magnitude of differences in licking observed between mouse DREADD groups and sexes. Parametric statistical tests, such as ANOVA, were conducted using R (Team, 2022). ANOVA effect size was gauged using partial eta squared (η^2^). Yuen’s two-sample trimmed mean *t*-test (WRS2 package in R; Mair and Wilcox, 2020) was applied for robust analysis of differences between two groups, including simple and main effects. Statistical effect size in Yuen’s test was gauged by ξ^, where values of 0.10, 0.30, and 0.50 correspond to small, medium, and large effects, respectively (Mair and Wilcox, 2020).

Estimation analyses were caried out using Gardner-Altman plots (Gardner and Altman, 1986) custom coded in MATLAB (release 2023b, MathWorks, Natick, MA). Here, all data points for each of two groups were displayed alongside their 20% trimmed mean and its 95% confidence interval. The 20% trimmed mean accommodated outliers by dropping the lower 20% and upper 20% of data points prior to averaging. Next, a bootstrap approach estimated the 95% confidence interval of the mean difference between the groups (i.e., the effect size in lick response units). To do this, each data group was randomly resampled with replacement, with the number of resampled data points equal to the group sample size, and the 20% trimmed mean computed. The difference between the resampled group means was stored, with this process carried out 1,000 times. Resampled differences were then sorted in ascending order to identify the 25^th^ (2.5%) and 975^th^ (97.5%) entries, which defined the 95% confidence interval range for the mean difference between groups. This calculation was repeated 100 times, with the average 95% confidence interval range reported in the results. The margin of error for the mean difference was half of the average confidence interval. The expected sampling error for the mean difference was represented by a probability distribution in Gardner-Altman plots. Estimation statistics allowed for visualization and interpretation of the effect size difference between two groups in the context of effect probability, data unit of measurement, and error/group variances.

## Results

We surgically prepared and tested 40 mCherry (19 females, 21 males) and 48 hM4Di (28 females, 20 males) mice in these studies. We analyzed behavioral data from mice that, following postmortem microscopy of brain tissue, showed bilateral cellular expression of mCherry in the external lateral region of the PB nucleus (**Figure 1**), which houses a dense cluster of *Calca* neurons (Huang et al., 2021). This pattern appeared in 59 mice (67% of all examined) and was taken as evidence of successful bilateral viral transfection of PB-*Calca* neurons with fluorescence control (mCherry) or inhibitory DREADD (hM4Di:mCherry) elements. Mice found to show only unliteral (*n* = 21, 24%) or no (*n* = 8, 9%) expression of mCherry in the external lateral PB area were not included in analyses.

**Figure 1.**
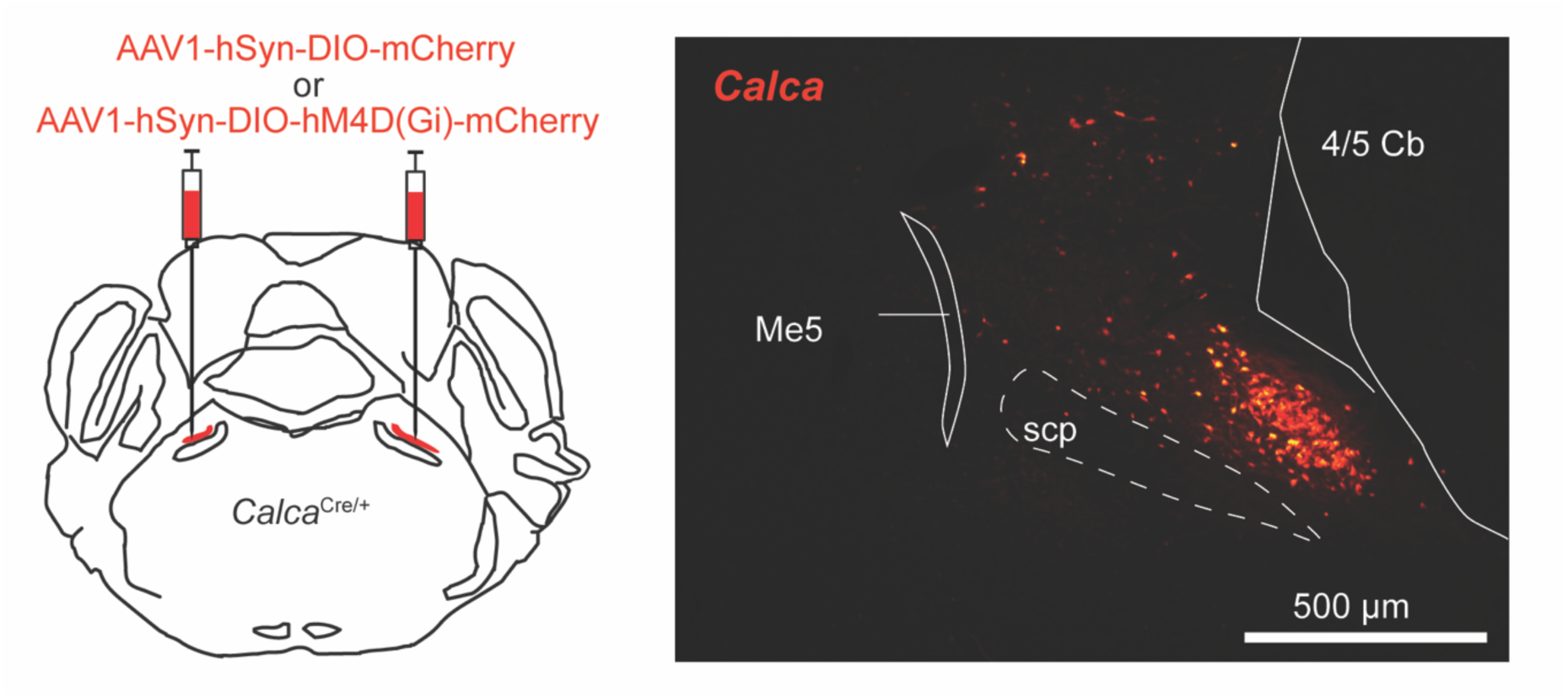
Left: Schematic of bilateral intracranial microinjections of Cre-dependent viruses encoding either mCherry or hM4Di:mCherry to the PB area in *Calca*^Cre/+^ mice. Right: Microscope image of brain tissue showing mCherry labeling of neurons in the external lateral parabrachial area in a *Calca*^Cre/+^ mouse. Abbreviations: Me5, mesencephalic trigeminal nucleus; scp, superior cerebellar peduncle; 4/5 Cb, cerebellar lobule.

### Viral transfection of PB-*Calca* neurons does not affect oromotor responding to water

We examined if the expression of exogenous fluorescence and DREADD proteins in *Calca* neurons disrupted normal licking behavior in mice. We observed that licks emitted to water during the first 5 trials of brief-access training without administration of CNO did not differ between mCherry (*n* = 28) and hM4Di (*n* = 31) mice (n.s. Yuen’s *t*-test, p = 0.308; **Figure 2**). Moreover, licks to water during brief-access training did not differ between female (*n* = 35) and male (*n* = 24) mice (n.s. Yuen’s *t*-test, p = 0.244).

**Figure 2.**
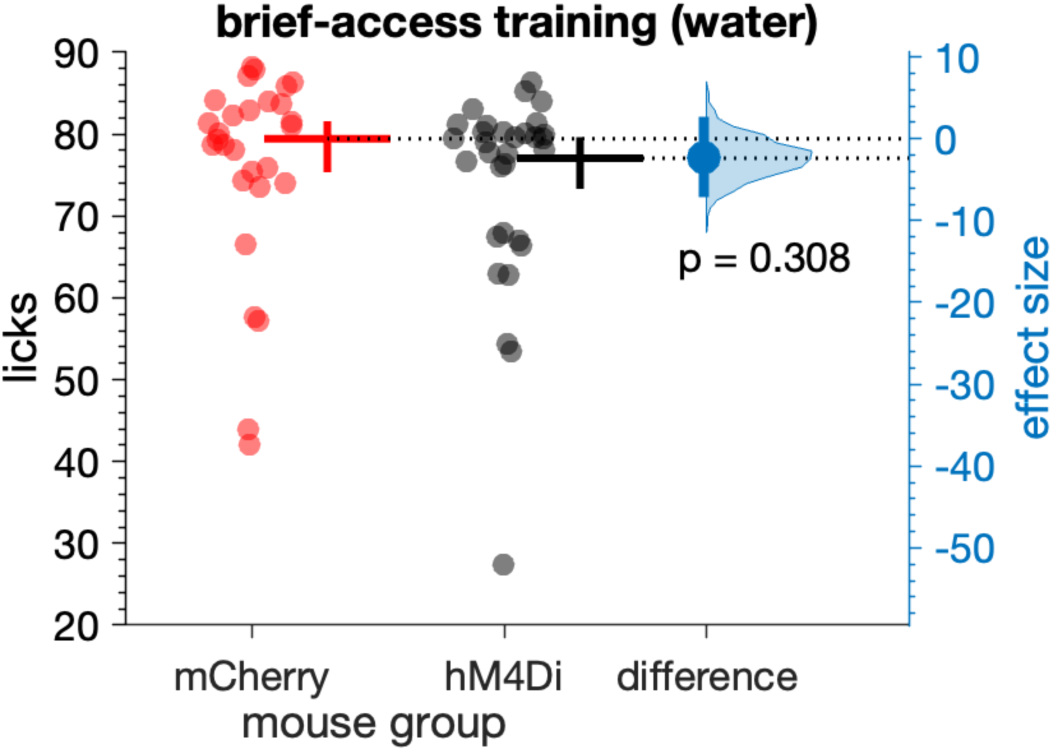
Licks emitted during brief-access training sessions with water did not differ (p = 0.308) between mCherry (*n* = 28) and hM4Di (*n* = 31) mice (markers). The mean (20% trimmed) number of licks (horizontal bar) and its 95% confidence interval (vertical bar) is plotted to the right of each distribution. The difference (in licks) between the sample means (effect size: blue circle) is shown to the right of the plot together with its 95% confidence interval (vertical blue bar).

On average, mCherry and hM4Di mice respectively made 79.5 and 77.1 licks during 10 sec brief-access training trials with water, without CNO (**Figure 2**). These lick rates approached the number of licks that B6 mice, which are a genetic background for *Calca*^Cre/+^ mice, could expectedly make with constant licking during a 10-sec fluid exposure trial with room temperature water (about 81 licks), as estimated from their established peak inter-lick interval (∼124 msec) (Boughter et al., 2007). Thus, adenoviral transfection of PB-*Calca* neurons did not influence normal oromotor responding in the absence of CNO.

### CNO does not affect preferences to innocuous oral temperatures in *Calca*;hM4Di mice

Our prior data show that when given a choice, thirsty mice prefer to lick water at a mild cool (15°C) rather than innocuous warm (35°C) temperature in brief-access tests conducted with temperature-controlled fluids (Li et al., 2024). Here, we examined if chemogenetic dampening of PB-*Calca* neurons would affect this behavior to gauge their role in preferences for innocuous orosensory stimuli.

A two-way ANOVA applied to data from 5 mCherry (3 females, 2 males) and 6 hM4Di (2 females, 4 males) mice undergoing water restriction revealed that 15°C water evoked more licks than 35°C water under CNO (main effect of temperature: *F*_1,9_ = 22.98, p = 0.00098, partial η^2^ = 0.719; **Figure 3**). This preference for mild cool water over warm agrees with our prior results (Li et al., 2024) and did not differ between mCherry and hM4Di mice (n.s. mouse group × temperature interaction, p = 0.734). Thus, PB-*Calca* neurons do not affect licking preferences towards innocuous orosensory cues, at least in the context of the thermal stimuli tested here.

**Figure 3.**
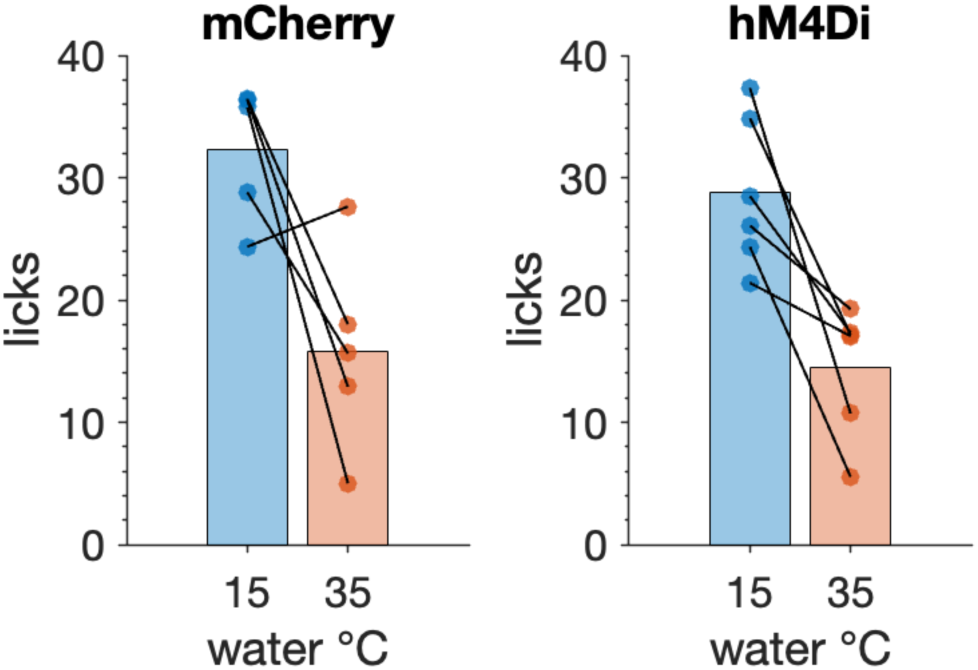
Both mCherry (*n* = 5) and hM4Di (*n* = 6) mice licked more to 15°C (mild cool) than 35°C (innocuous warm) water during brief-access fluid exposure tests conducted under CNO (p = 0.00098). Lines connect responses (markers) made by one mouse. Bars are 10% trimmed means.

### CNO variably reduces bitter taste avoidance in *Calca*;hM4Di mice

#### Quinine

Orosensory responses to the bitter taste stimulus quinine were compared between 21 mCherry (12 females, 9 males) and 25 hM4Di (16 females, 9 males) water-restricted mice. We found that while under CNO, mCherry and hM4Di mice made similar licks to water (0 mM) offered with quinine solutions during brief-access tests (n.s. independent samples *t*-test, p = 0.104; **Figure 4A**). A three-way ANOVA revealed that adulterating water with quinine decreased licking in both mouse groups as quinine concentration increased (main effect of quinine concentration, *F*_3,126_ = 212, p < 0.001, partial η^2^ = 0.835; **Figure 4A**). Yet the degree of this effect differed between mCherry and hM4Di mice (mouse group × quinine concentration interaction, *F*_3,126_ = 4.08, p = 0.0084, partial η^2^ = 0.089), with hM4Di mice appearing to show, on average, less of a reduction in responding (i.e., greater licks to quinine) while under CNO.

**Figure 4.**
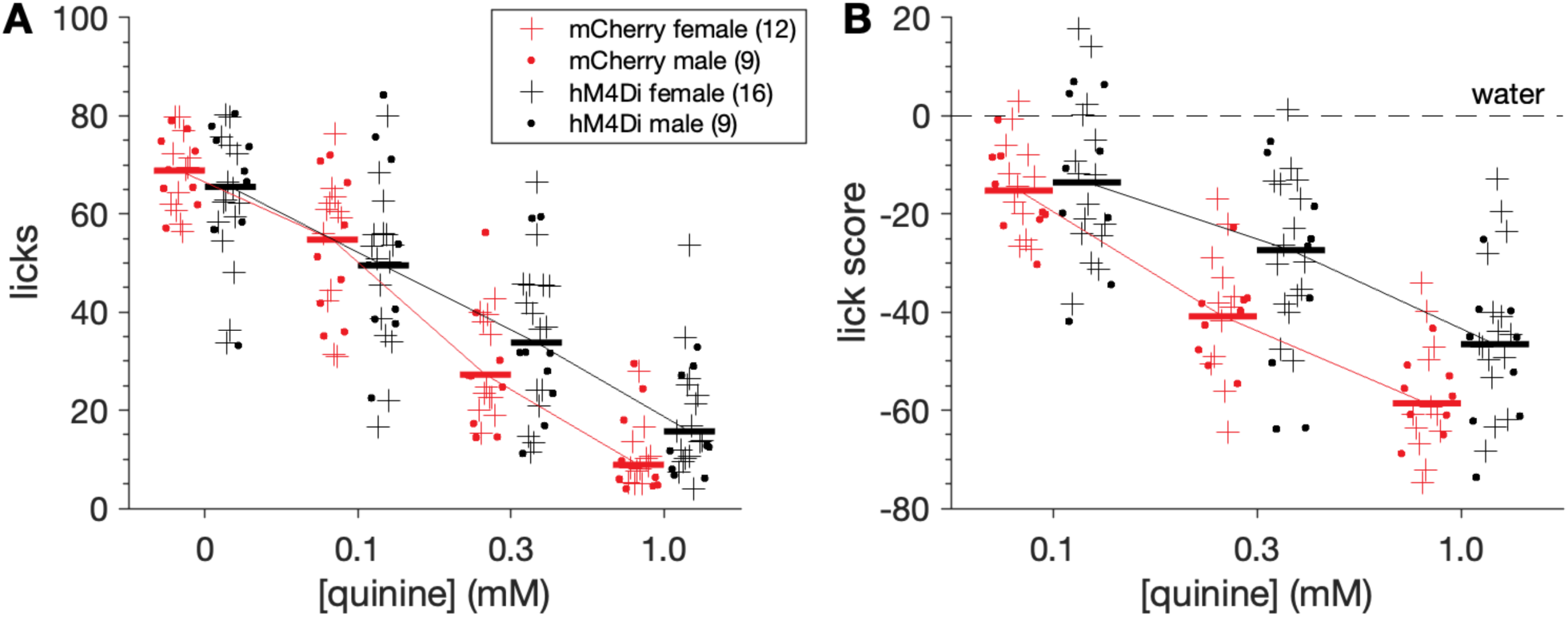
hM4Di mice show reduced orosensory avoidance of quinine while under CNO. **A**, Concentration-response functions showing the number of licks to water (0 mM) and quinine solutions (0.1 to 1.0 mM) for all mice (markers) in both mouse groups and sexes (legend). **B**, Quinine concentration-response functions re-plotted using licks scores, where points represent the number of stimulus licks minus water licks for all mice. Horizontal bars in both panels are 20% trimmed means. Factorial analyses revealed that stimulus concentration and mouse line interacted to influence licks (p = 0.0084) and lick scores (p = 0.044) to quinine.

Analysis of lick scores, which standardized responses as reductions in licks from water licking, revealed that lick differences to quinine between mCherry and hM4Di mice were conditioned on stimulus concentration (three-way ANOVA: mouse group × quinine concentration interaction, *F*_2,84_ = 3.26, p = 0.044, partial η^2^ = 0.072; **Figure 4B**). Robust simple effects tests found that while under CNO, lick scores to 0.1 mM quinine did not differ between mouse groups (n.s. Yuen’s *t*-test, p = 0.701; **Figure 5A**). However, 0.3 mM quinine evoked higher lick scores in hM4Di mice (Yuen’s *t*-test, *t*_23.6_ = 3.01, p = 0.006, ξ^= 0.55). On average, hM4Di mice emitted about 14 more licks to 0.3 mM quinine (a 33% increase) than mCherry mice during 10-sec trials, with a 95% confidence interval for this difference of 3 to 22 more licks (**Figure 5B**). The margin of error for this increase was about 9 licks, which was moderately lower than the effect size of 14 licks.

**Figure 5.**
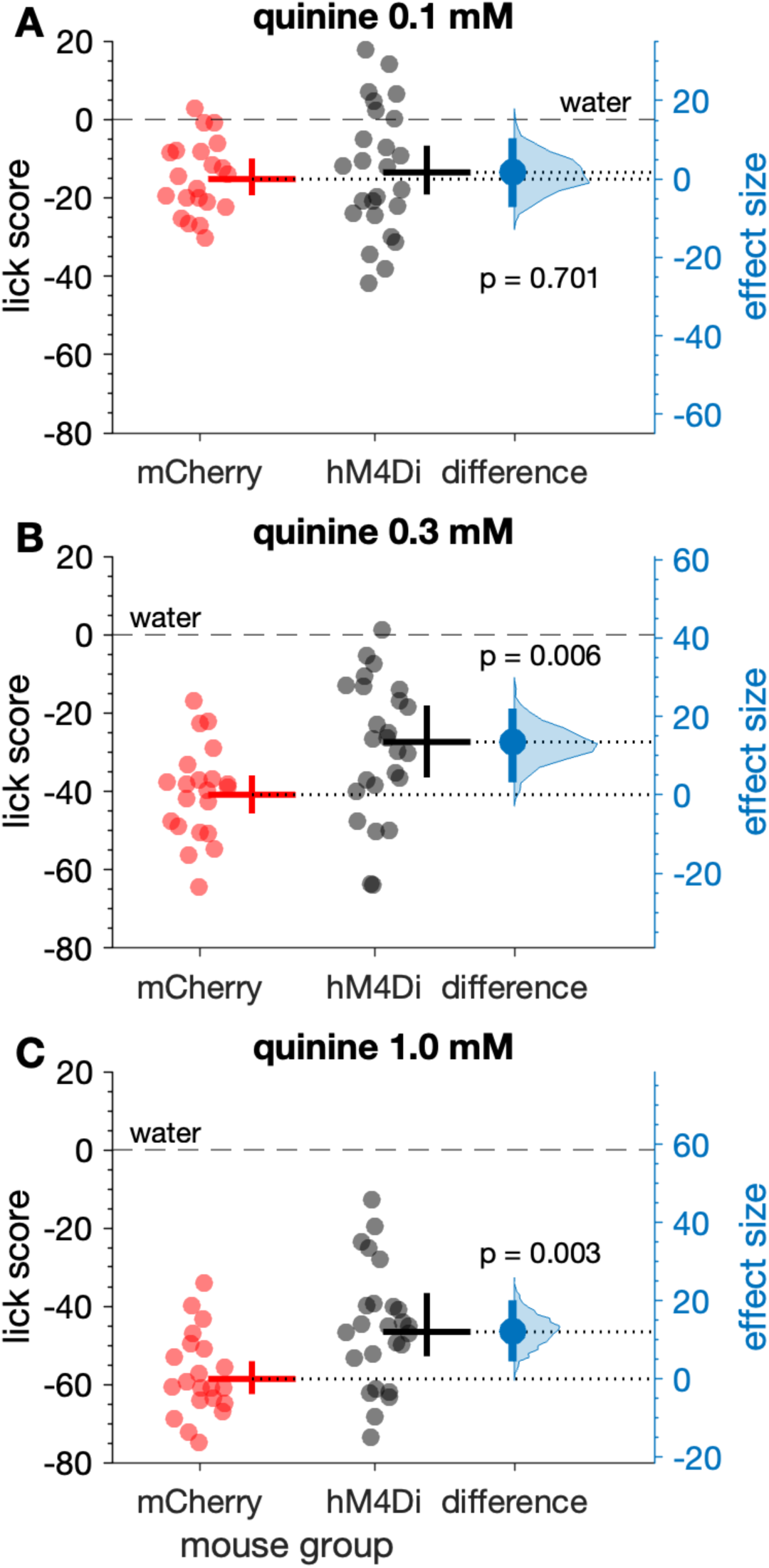
Lick scores to elevated concentrations of quinine were larger in hM4Di (*n* = 25) compared to mCherry (*n* = 21) mice (markers), all receiving CNO. Lick scores to 0.1 mM quinine did not differ between mCherry and hM4Di mice (**A**, p = 0.701). In contrast, lick scores to 0.3 mM (**B**, p = 0.006) and 1.0 mM (**C**, p = 0.003) quinine were higher in hM4Di mice under CNO. For each panel, the mean (20% trimmed) number of licks (horizontal bar) and its 95% confidence interval (vertical bar) is plotted to the right of each distribution. The difference between the sample means (effect size: blue circle) is shown to the right of the plot together with its 95% confidence interval (vertical blue bar).

Moreover, lick scores to 1.0 mM quinine were higher in hM4Di compared to mCherry mice (Yuen’s *t*-test, *t*_25.2_ = 3.29, p = 0.003, ξ^= 0.60), with hM4Di mice making about 12 more mean licks (a 21% increase) in 10-sec trials compared to mCherry controls (**Figure 5C**). The 95% confidence interval for this increase was 4 to 20 more licks (margin of error = 8 licks).

Although some of the largest increases in responding to quinine observed in hM4Di mice emerged in females (**Figure 4A, 4B**), sex did not significantly interact with mouse group and quinine concentration to influence quinine lick scores under CNO (n.s. three-way interaction, p = 0.582; n.s. sex × mouse group interaction, p = 0.661). Finally, latency to first lick was not influenced by mouse group or quinine concentration (n.s. mouse group × concentration interaction, p = 0.704; n.s. effect of mouse group, p = 0.326; n.s. effect of concentration, p = 0.119), which suggested olfactory/vapor cues did not affect responses to quinine.

#### Cycloheximide

Orosensory responses to the bitter taste stimulus cycloheximide were examined in a different cohort of 5 mCherry (3 females, 2 males) and 6 hM4Di (3 females, 3 males) water-restricted mice. Under CNO, mCherry and hM4Di mice showed similar licks to water offered with cycloheximide solutions during tests (n.s. independent samples *t*-test, p = 0.330; **Figure 6A**). Both mouse groups showed concentration-dependent reductions in licks (two-way ANOVA: main effect of concentration, *F*_3,27_ = 74.21, p < 0.001, partial η^2^ = 0.892; **Figure 6A**) and lick scores (two-way ANOVA: main effect of concentration, *F*_2,18_ = 61.85, p < 0.001, partial η^2^ = 0.873; **Figure 6B**) to cycloheximide. Yet unlike quinine, these reductions were similar between groups (n.s. mouse group × concentration interaction on licks scores, p = 0.974). Relatedly, mean lick scores to cycloheximide collapsed across concentration did not differ between hM4Di and mCherry mice under CNO (n.s. Yuen’s *t*-test, p = 0.327; **Figure 6C**). Underpowered sample sizes precluded analyses of cycloheximide data by sex, albeit no observable differences between females and males appeared in plotted data (**Figure 6A, 6B**).

**Figure 6.**
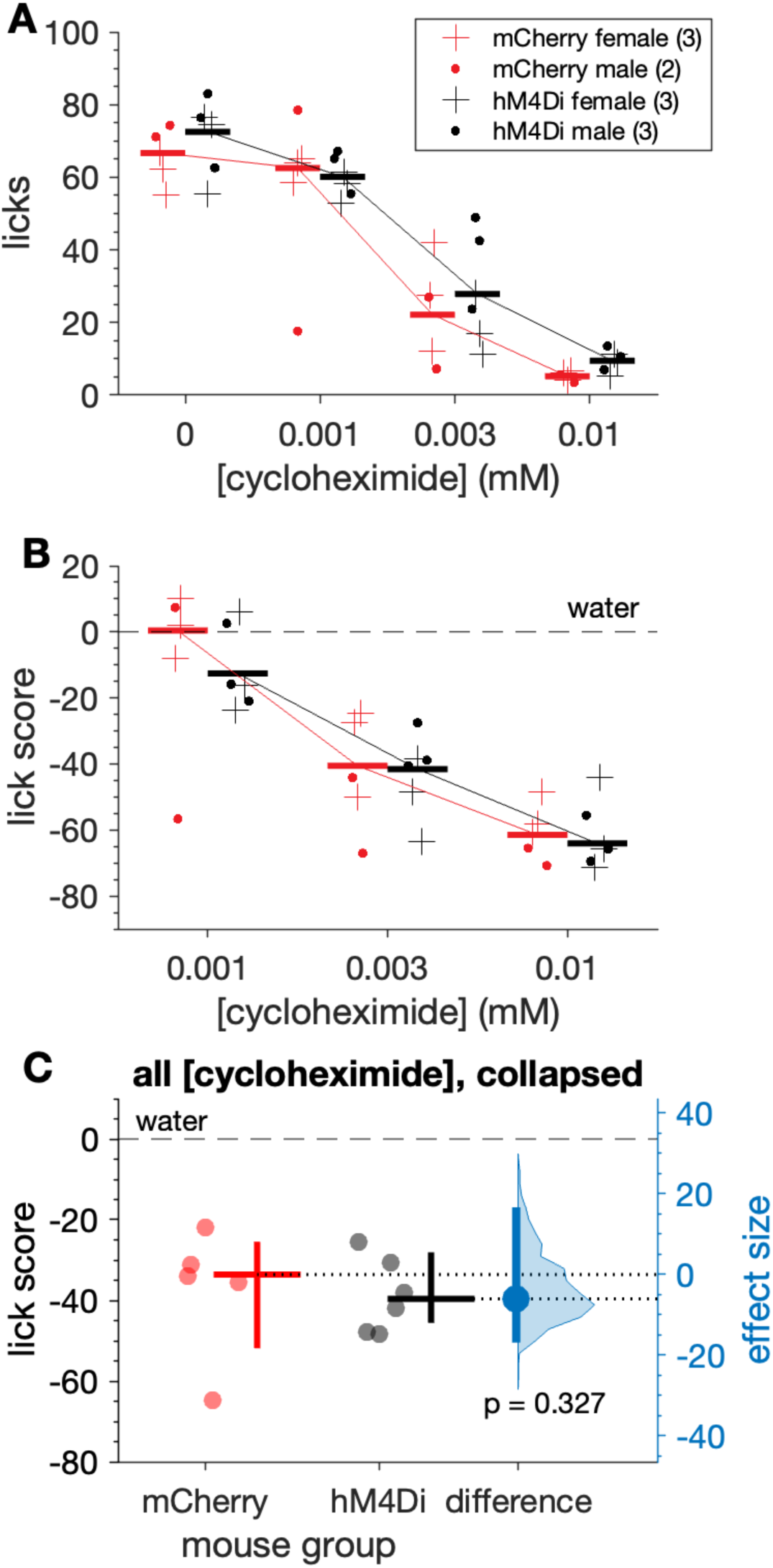
Orosensory avoidance of cycloheximide did not differ between hM4Di and mCherry mice under CNO. Plots show concentration-response functions for cycloheximide licks (**A**) and lick scores (**B**) by hM4Di (*n* = 6) and mCherry (*n* = 5) mice (markers) administered CNO. Horizontal bars are 20% trimmed means. **C**, Lick scores to cycloheximide collapsed across concentration did not differ (p = 0.327) between mouse groups. The mean (20% trimmed) number of licks (horizontal bar) and its 95% confidence interval (vertical bar) is plotted to the right of each distribution. The difference between the sample means (effect size: blue circle) is shown to the right of the plot together with its 95% confidence interval (vertical blue bar).

Altogether, these results indicate that CNO variably suppressed orosensory avoidance of bitter taste stimuli in mice where the inhibitory DREADD hM4Di was expressed in PB-*Calca* neurons. In these (hM4Di) mice, CNO significantly increased licking to quinine solutions with some variance in this increase noted across animals. Based on this variance, a range of only marginal to larger increases in responding to quinine under CNO were compatible with our data. Quinine behavioral response variance was not linked to sex and is further discussed below. Unlike quinine, orosensory responses to cycloheximide were not significantly influenced by CNO. This may reflect participation of PB-*Calca* neurons in known functional differences between cycloheximide and quinine.

### CNO causes sex-dependent reductions in sucrose taste preference in *Calca*;hM4Di mice

We analyzed orosensory responses to solutions of a preferred sugar, sucrose, collected from 17 mCherry (11 females, 6 males) and 26 hM4Di (16 females, 10 males) mice. Both mCherry and hM4Di mice emitted increased licks to sucrose at elevated stimulus concentrations during the sucrose training sessions conducted under water restriction conditions, prior to CNO tests (three-way ANOVA: main effect of concentration, *F*_4,156_ = 6.14, p = 0.00013, partial η^2^ = 0.136; **Figure 7**). These increases were similar between mouse groups (n.s. mouse group × sucrose concentration interaction, p = 0.794) and sexes (n.s. sex × mouse group × sucrose concentration interaction, p = 0.620; n.s. sex × mouse group interaction, p = 0.278).

**Figure 7.**
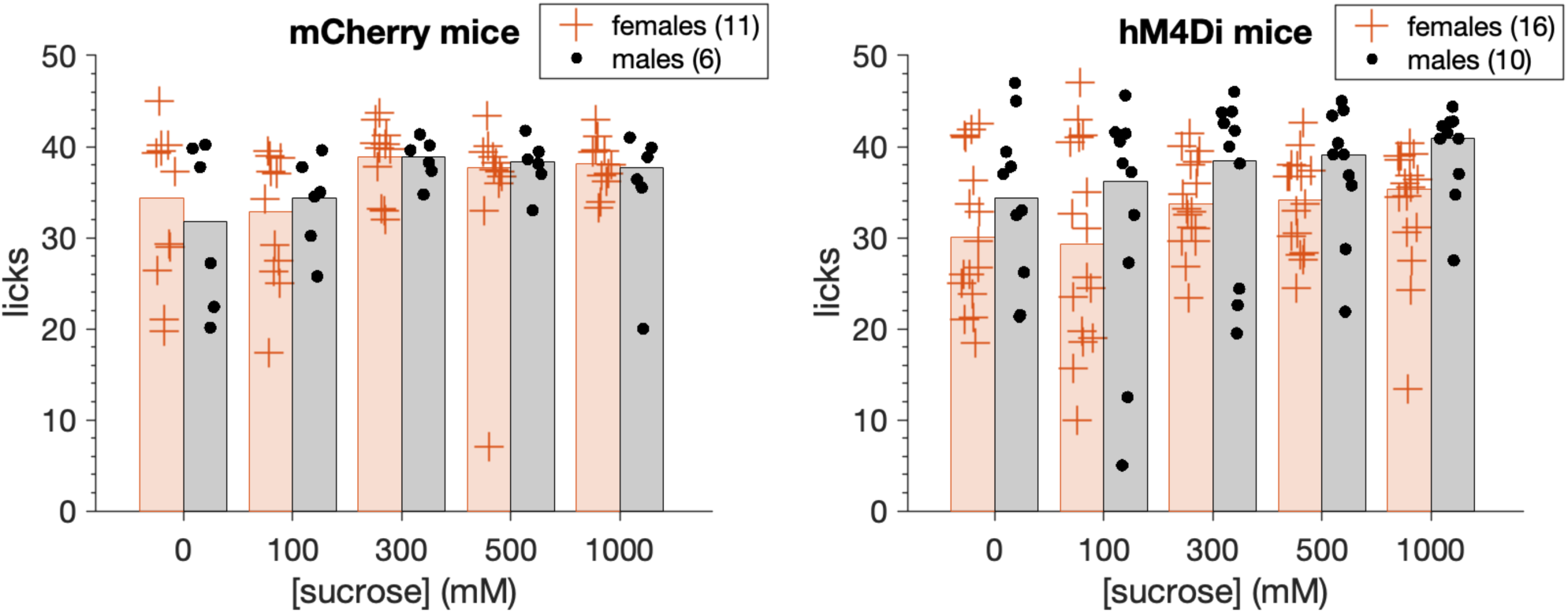
Water-restricted mCherry (*n* = 17) and hM4Di (*n* = 26) mice (markers) increased (p = 0.00013) their licks emitted to sucrose solutions at elevated stimulus concentrations during pre-CNO sucrose training sessions. These elevations were similar between mouse groups (p = 0.794) and sexes (p = 0.620). Bars are 20% trimmed means.

Following sucrose training, mCherry and hM4Di mice entered brief-access tests with the sucrose series while water replete (not thirsty) and after receiving CNO. A three-way ANOVA revealed that in the absence of thirst drive, both mouse groups showed concentration-dependent increases in licks to sucrose when tested under CNO (main effect of concentration, *F*_3,117_ = 77.72, p < 0.0001, partial η^2^ = 0.666; **Figure 8A, 8B**). However, the magnitude of these increases differed between mCherry and hM4Di mice in a manner that was conditioned on sex (sex × mouse group × sucrose concentration interaction, *F*_3,117_ = 3.15, p = 0.027, partial η^2^ = 0.075). Inspection of plotted data revealed that orosensory responses to sucrose were markedly and selectively impaired in male hM4Di mice, which displayed, on average, fewer licks to elevated sucrose concentrations than female hM4Di, and all mCherry, mice (**Figure 8B, 8A**).

**Figure 8.**
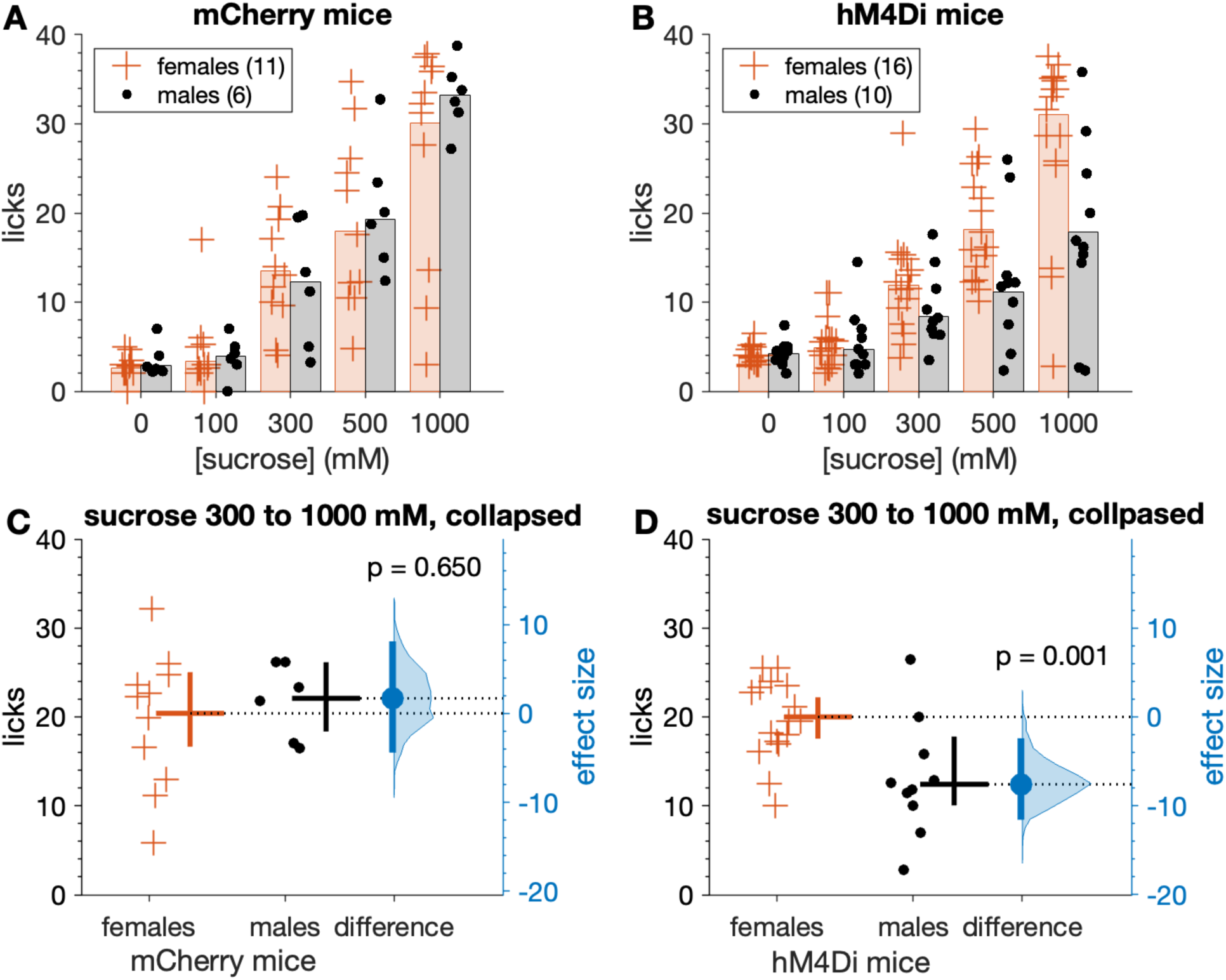
Male hM4Di mice show reduced licks to sucrose under CNO when water replete (not thirsty). Top row shows the number of licks emitted across the sucrose series for mCherry (**A**) and hM4Di (**B**) mice (markers). Bars are 20% trimmed means. When collapsed across the three highest (most salient) concentrations, mean licks to sucrose did not differ between female and male mCherry mice (**C**, p = 0.650). However, male hM4Di mice made fewer licks to sucrose than female hM4Di mice (**D**, p = 0.001). Female hM4Di mice responded similarly to mCherry control mice (see text). In panels C and D, the mean (20% trimmed) number of licks (horizontal bar) and its 95% confidence interval (vertical bar) is plotted to the right of each distribution. The difference (in licks) between the sample means (effect size: blue circle) is shown to the right of the plot together with its 95% confidence interval (vertical blue bar).

When data were collapsed across the three highest (most salient) sucrose concentrations tested (300, 500, and 1000 mM), male hM4Di mice emitted, on average, about 8 fewer licks to sucrose (a 38% decrease) during 5-sec exposure trials compared to female hM4Di mice when all mice were administered CNO (Yuen’s *t*-test, *t*_11.8_ = 4.31, *p* = 0.001, ξ^= 0.78; **Figure 8D**). The 95% confidence interval for this decrease was about 2 to 12 less licks, with a margin of error of 5 fewer licks. In contrast, licks to salient concentrations of sucrose under CNO did not differ between female and male mCherry mice (n.s. Yuen’s *t*-test, p = 0.65; **Figure 8C**), which both licked sucrose at the same rate as female hM4Di mice (n.s. Yuen’s *t*-test, p = 0.496).

The same trend emerged when brief-access licking responses were standardized for each mouse using lick ratios, where their lick count to each sucrose concentration was divided by an estimate of their maximal potential licking rate. Lick ratios to sucrose captured under CNO increased with elevations in sucrose concentration in a mouse group- and sex-dependent manner, with male hM4Di mice displaying a unique reduction in lick ratio responses compared to the other mouse groups (three-way ANOVA, sex × mouse group × sucrose concentration interaction, *F*_3,117_ = 3.29, p = 0.023, partial η^2^ = 0.078; **Figure 9A, 9B**). Specifically, lick ratios to 300, 500, and 1000 mM sucrose (collapsed) were 39% lower in male hM4Di compared to female hM4Di mice under CNO (Yuen’s *t*-test, *t*_12.6_ = 5.06, p = 0.00024, ξ^= 0.75; **Figure 9D**). The 95% confidence interval for this reduction ranged from 11% to 58% lower (margin of error = 23%). On the other hand, lick ratios collapsed across 300, 500, and 1000 mM sucrose did not differ between female and male mCherry mice (n.s. Yuen’s *t*-test, p = 0.525; **Figure 9C**).

**Figure 9.**
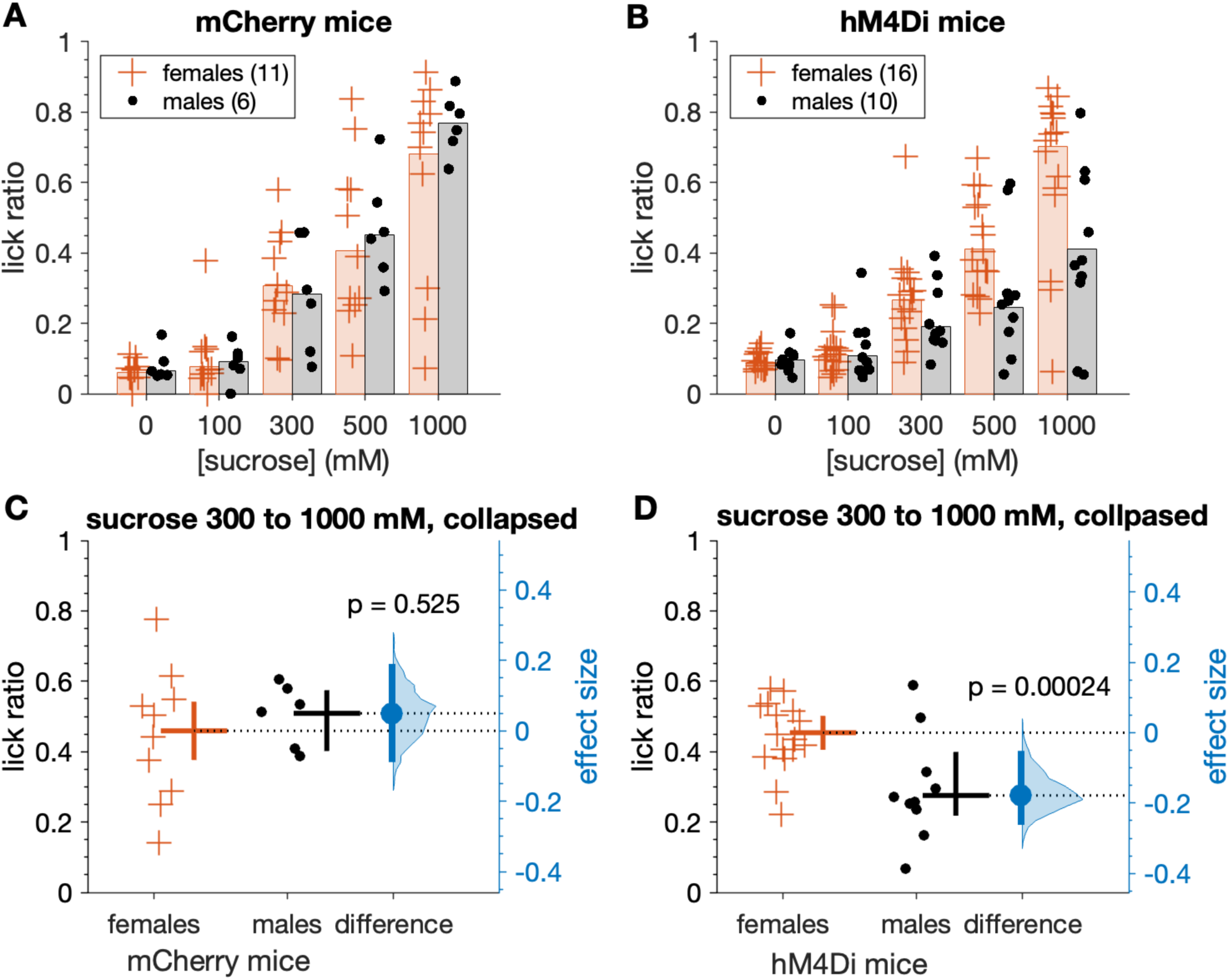
Male hM4Di mice show reduced lick ratios to sucrose under CNO when water replete. **A**, Sucrose lick ratios for mCherry mice (markers). Bars are 20% trimmed means. **B**, same as A but for hM4Di mice. When collapsed across the three highest (most salient) concentrations, mean lick ratios to sucrose did not differ between female and male mCherry mice (**C**, p = 0.525). In contrast, mean sucrose lick ratios were reduced in male compared to female hM4Di mice (**D**, p = 0.00024). In panels C and D, the mean (20% trimmed) lick ratio (horizontal bar) and its 95% confidence interval (vertical bar) is plotted to the right of each distribution. The difference between the sample means (effect size: blue circle) is shown to the right of the plot together with its 95% confidence interval (vertical blue bar).

The lick ratio standardization used here is appropriate to gauge orosensory avidity to appetitive taste stimuli that stimulate licking, like sucrose (Glendinning et al., 2002; Dotson and Spector, 2004; Ellingson et al., 2009). Notably, many mCherry and female hM4Di mice displayed lick ratios that approached 1 (maximal licking) for the most salient sucrose concentrations tested, albeit most male hM4Di mice showed markedly reduced ratio responding (**Figure 9A, 9B**).

Finaly, latency to first lick to sucrose solutions did not differ across concentrations (three-way ANOVA, n.s. effect of sucrose concentration, p = 0.834) and was not influenced by mouse group (n.s. mouse group × concentration interaction, p = 0.656; n.s. effect of group, p = 0.415) or sex (n.s. sex × mouse group × concentration interaction, p = 0.999). Thus, olfactory/vapor cues did not impact orobehavioral responses to sucrose.

Altogether, these and the above data reveal that CNO suppressed hedonic licking of sucrose selectively in male hM4Di mice, where the inhibitory DREADD hM4Di was expressed in PB-*Calca* neurons. This trend agrees with a recently described ability of *Calca* neurons to signal aversive *and* appetitive stimuli (Kim et al., 2024a), as below. The current results suggest that *Calca* neuron participation in appetitive taste depends on sex.

A subset of the female (*n* = 9) and male (*n* = 7) hM4Di mice examined for sucrose preferences were also tested in a brief-access setting with sucrose performed without daily administration of CNO. In these tests, no significant difference in responding between sexes was found in the absence of CNO-activated neuronal dampening via hM4Di (n.s. Yuen’s *t*-test, p = 0.095; **Figure 10**). This suggested that differences in responding to sucrose between female and male hM4Di mice (**Figure 8, 9**) relied on CNO, which supported temporary suppression of PB-*Calca* cells in these mice. Nevertheless, inspection of the plotted data and sex difference confidence interval for the no-CNO tests showed that sucrose responses by males did not fully match female levels, with continued reductions in male responding compatible with our data (**Figure 10**). We discuss this below.

**Figure 10.**
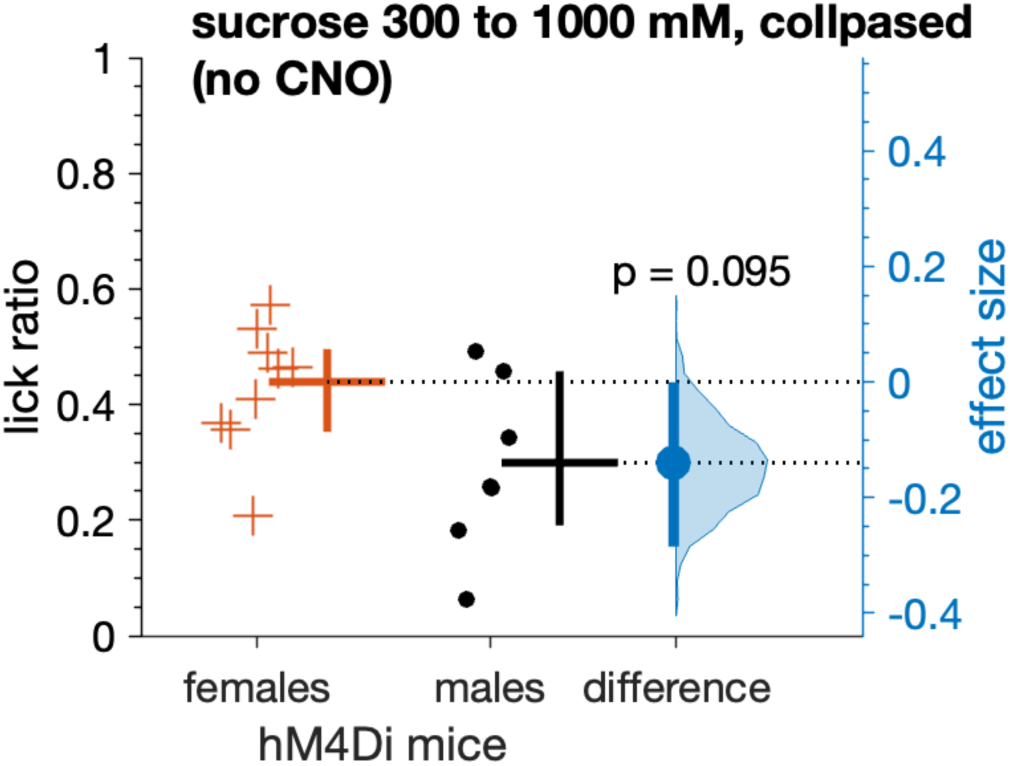
Without CNO, lick ratios to sucrose, collapsed across the three highest concentrations, did not differ between female and male hM4Di mice (p = 0.095). The mean (20% trimmed) lick ratio (horizontal bar) and its 95% confidence interval (vertical bar) is plotted to the right of each distribution. The difference between the sample means (effect size: blue circle) is shown to the right of the plot together with its 95% confidence interval (vertical blue bar).

## Discussion

The present study provides evidence that PB-*Calca* neurons participate in mouse orosensory responses to hedonically diverse taste stimuli. Using brief-access fluid exposure tests, we found that chemogenetic dampening of activity targeted to PB-*Calca* neurons increased mouse licking acceptance of the bitter taste stimulus quinine, with observed variability in the magnitude of this effect across mice. What is more, chemogenetic dampening of PB-*Calca* neurons decreased mouse licking preference for the appetitive sugar (sweet) sucrose. The latter result was conditioned on sex, with the reduced responding to sucrose that followed administration of CNO appearing selectively in male hM4Di mice. During each daily test session, mice were able to lick each taste stimulus concentration for up to a maximum of 40 (sucrose) or 50 (bitter stimuli) sec. Thus, the noted effects arose during brief encounters with these stimuli, reflective of sensory-guided responses.

This dual effect on aversive (quinine) and preferred (sucrose) tastes may appear perplexing in the context that PB-*Calca* neurons were established to have roles in protective responses to unfavorable and noxious conditions (e.g, Carter et al., 2013; Campos et al., 2018; Kang et al., 2022). Selective toxin-induced inactivation of PB-*Calca* neurons was also reported to have no effect on mouse licking preferences to the non-caloric sweetener saccharin (Jarvie et al., 2021). However, recent data have implied these neurons display a more diverse response repertoire that incudes activation to appetitive stimuli (Kim et al., 2024b). Notably, PB-*Calca* neurons appear to respond to preferred and aversive stimuli by changes in response frequency and use of frequency-modulated neurotransmission. In mice, drinking sucrose (appetitive) and tail pinch (aversive) caused low and high rates of spiking in PB-*Calca* neurons that associated with glutamate or neuropeptide release, respectively, which may contribute different component features to sensations (Kim et al., 2024a). The present results suggest that participation of PB-*Calca* neurons in appetitive taste-guided behavior is critically linked to sex, with normal functioning in PB-*Calca* cells needed for male, but not female, mice to express typical licking preference responses to sucrose. Prior studies that examined how PB-*Calca* neurons may affect mouse fluid licking associated with taste did not address or report sex effects.

The present data also imply that PB-*Calca* neurons can exert a variable influence on avoidance behavior across bitter taste stimuli. The increased acceptance of quinine shown, on average, by hM4Di mice on receipt of CNO was dependent on stimulus intensity, with these mice emitting more licks to quinine at elevated (≥0.3 mM) but not reduced (0.1 mM) concentrations compared to mCherry controls. In contrast, hM4Di and mCherry mice administered CNO did not differ in their licking responses to the bitter tastant cycloheximide. This differential effect of CNO on quinine and cycloheximide avoidance in hM4Di mice associates with established functional differences between these stimuli. In rodents, oral presence of quinine normally evokes robust electrophysiological responses in cranial nerve (CN) VII and CN IX (Dahl et al., 1997; Inoue et al., 2001; Danilova and Hellekant, 2003; Damak et al., 2006), which respectively supply rostral and caudal lingual taste bud fields and are evidenced to support different aspects of taste processing and behavior under certain conditions (Travers et al., 1987; Spector and Grill, 1992; Spector et al., 1997; St. John and Spector, 1998; see also St John and Boughter, 2004). In contrast to quinine, cycloheximide induces strong gustatory activity mainly in CN IX (Danilova and Hellekant, 2003; Damak et al., 2006; Hettinger et al., 2007). Differences in gustatory neural responses to quinine and cycloheximide also emerge in the rodent CNS, where these stimuli evoke partly overlapping but distinct neural population responses in brain stem taste structures (Geran and Travers, 2006; Wilson et al., 2012), including the PB nucleus (Geran and Travers, 2009).

Speculatively, PB-Calca neurons could participate in circuits that support central distinctions in neural information between quinine and cycloheximide given that perturbation of these neurons presently had different results on orosensory guided behaviors to these bitters. Additional PB neural types are likely also involved with these circuits, including Satb2-positive neurons found to contribute to quinine licking avoidance behaviors alongside *Calca* cells (Jarvie et al., 2021). Moreover, oral presence of cycloheximide stimulates taste-active neurons found in PB areas (Geran and Travers, 2009; Li and Lemon, 2019; Li et al., 2022) populated by *Calca* neurons (Huang et al., 2021). When coupled with the present findings, this result opens the possibility that cycloheximide taste does engage PB-*Calca* neurons but that actions of another neural class or circuit may exert greater influence on taste reactions to cycloheximide. Future studies on cell types supporting orobehavioral responses to other functionally diverse bitters may shed deeper light on this hypothesis and the significance of heterogeneity among bitter taste responses.

In the present study, mice displayed notable variance in responding to select taste stimuli following receipt of CNO. For example, while the mean lick score to 0.3 mM quinine was significantly higher in hM4Di mice, a few of these animals showed no or only a mild increase in licks compared to the average mCherry control group response. In contrast, other hM4Di mice displayed substantially elevated responding, licking 0.3 mM quinine at water-like levels under CNO (**Figure 5B**). The confidence interval for the difference between mouse groups demonstrated that a range of relatively small to larger increases in quinine licking by hM4Di mice are compatible with our data and could arise on replication (**Figure 5B**).

It is important to consider that viral-based delivery of DREADDs to neural tissues may not affect all neurons of a targeted population (Smith et al., 2016). Within this context, DREADD effects on activity in transfected neurons are likely not complete such that CNO activation of hM4Di may only induce modest hyperpolarization and dampen, but not fully silence, the cellular response (Roth, 2016; Smith et al., 2016). Variance in these parameters across hM4Di mice could have contributed to variance in the present behavioral effects that followed chemogenetic perturbations. Yet mCherry mice showed reduced but marked variance in licking responses in some cases (e.g., **Figure 5B**), suggesting some of the present behavioral variance may reflect latent factors not examined here. Nevertheless, we aimed to test sizable numbers of experimental and control mice to accommodate variance. Notably, all mice included in analyses showed at least bilateral mCherry labeling of neurons, indicative of expression of hM4Di:mCherry or mCherry alone, in the external lateral PB nucleus, which is densely populated by *Calca* cells (Huang et al., 2021). Fewer mice were included in analyses of cycloheximide data because not all tested met this criterion.

We observed that the mean licking response to sucrose did not significantly differ between female and male hM4Di mice in the *absence* of CNO (**Figure 10**). This suggests that the reduced sucrose preference found in male hM4Di mice with CNO (**Figure 8, 9**) was conditioned on the DREADD influence on PB-*Calca* neurons. Nevertheless, plotting all data suggested that some individual male hM4Di mice may have responded less to sucrose during no-CNO tests, with the confidence interval of the difference between sexes indicating reduced male responses are plausible (**Figure 10**). Because most of these no-CNO data were collected after tests with CNO, this trend may speculatively reflect carryforward of reduced responding or learning phenomena. While additional work is needed to address how *Calca* neurons, and sex, influence this process, prior studies have implicated PB-*Calca* neurons with roles in ingestive learning (Carter et al., 2015; Chen et al., 2018).

The present data support a novel role for PB-*Calca* neurons in taste-guided behaviors to aversive and appetitive gustatory stimuli, with appetitive orosensory responses affected by these cells displaying significant dependence on sex. While arising in the external lateral PB area, *Calca* cells have also been identified in other PB regions, including dorsal lateral, medial, and waist areas (Schwaber et al., 1988; Huang et al., 2021) closely associated with taste (Geran and Travers, 2009; Tokita et al., 2014; Tokita and Boughter, 2016). Future studies on *Calca* circuits originating from diverse PB locations may help further delineate their functional roles in gustation.

## Conffict of interest statement

The authors declare no competing interests, financial or otherwise.

## Acknowledgements

Research reported in this publication was supported by NIH grants DC011579 and DC020843 to C.H.L. The content is solely the responsibility of the authors and does not necessarily represent the official views of the NIH. Portions of these data were presented in abstract form at the 2022 C 2024 meetings of the Association for Chemoreception Sciences. Current Address for Md Ali: Department of Neural and Pain Sciences, University of Maryland School of Dentistry, Baltimore, MD, 21201. The authors thank Traci Redwine and Damilola Adesina for assistance with histology.

## Author contributions

Conception and design of research: C.H.L.

Performed experiments: J.L., M.S.S.A., N.M.N., K.T.Z., C.J.R.

Analyzed data: C.H.L.

Prepared figures: C.H.L., J.L.

Drafted manuscript: C.H.L.

Approved final version of manuscript: C.H.L., M.S.S.A., J.L., N.M.N., C.J.R., K.T.Z.

## Notes

### Competing Interest Statement

The authors have declared no competing interest.

